# Elevated FOXG1 supports exit from quiescence in neural stem cells through FoxO6

**DOI:** 10.1101/2022.06.30.498283

**Authors:** Kirsty M Ferguson, Carla Blin, Claudia Garcia-Diaz, Harry Bulstrode, Raul Bardini Bressan, Katrina McCarten, Steven M. Pollard

## Abstract

The molecular mechanisms controlling the balance of quiescence and proliferation in adult neural stem cells (NSCs) are often deregulated in brain cancers such as glioblastoma (GBM). Previously, we reported that FOXG1, a forebrain-restricted neurodevelopmental transcription factor, is frequently upregulated in glioblastoma stem cells (GSCs) and limits the effects of cytostatic pathways, in part by repression of the tumour suppressor *Foxo3*. Here, we show that increased FOXG1 upregulates *FoxO6*, a more recently discovered FoxO family member with potential oncogenic functions. Although genetic ablation of *FoxO6* in proliferating NSCs has no effect on the cell cycle or entry into quiescence, we find that *FoxO6*-null NSCs can no longer efficiently exit quiescence following FOXG1 elevation. Increased *FoxO6* results in the formation of large acidic vacuoles, reminiscent of Pak1-regulated macropinocytosis. Consistently, Pak1 expression is upregulated by FOXG1 overexpression and downregulated upon FoxO6 loss in proliferative NSCs. These data suggest a pro-oncogenic role for FoxO6 in controlling the exit from quiescence in NSCs, and shed light on the functions of this underexplored FoxO family member.

**Research highlights:**

- FoxO6 is a downstream effector of elevated FOXG1 in mouse NSCs and GSCs.
- Upregulation of FoxO6 is necessary for FOXG1 to drive efficient quiescence exit of NSCs.
- FoxO6 overexpression stimulates macropinocytosis, a process regulated by the actin cytoskeleton regulator Pak1.
- Pak1 is upregulated by FOXG1 overexpression and downregulated upon FoxO6 loss.

## Introduction

Stem cell fate is orchestrated by gene regulatory networks of ‘lineage-specific’ master regulatory transcription factors (Graf & Enver, 2009). Just as tissues rely on these factors for proper development, cancers can subvert developmental networks to impose a stem cell-like state that underpins tumour growth (Roy & Hebrok, 2015; Huilgol *et al*, 2019). Glioblastoma multiforme (GBM), the most common and aggressive primary adult brain cancer, is driven by glioblastoma stem cells (GSCs) that display neural stem cell (NSC) characteristics (Singh *et al*, 2003; Pollard *et al*, 2009; Richards *et al*, 2021). GSCs frequently overexpress key neurodevelopmental transcription factors to drive their self-renewal and restrict differentiation (Engström *et al*, 2012; Carén *et al*, 2015; Suva *et al*, 2014; Singh *et al*, 2017). One such factor is the Forkhead box transcription factor, FOXG1. FOXG1 has important roles in telencephalon development and *in vitro* reprogramming (Lujan *et al*, 2012; Xuan *et al*, 1995; Bulstrode *et al*, 2017). It is one of the most consistently overexpressed genes across GBM molecular subtypes, and high levels are associated with adverse outcomes (Engström *et al*, 2012; Robertson *et al*, 2015; Wang *et al*, 2018; Verginelli *et al*, 2013). Understanding the molecular mechanisms through which FOXG1 operates in NSCs and GSCs is therefore of great interest.

Both GSCs and genetically normal NSCs are known to be heterogeneous with regards to cell cycle status, with cells spanning a continuum from dormant to quiescent and proliferative states (non-cycling, slow-cycling and fast cycling, respectively) (Codega *et al*, 2014; Dulken *et al*, 2017; Marqués-Torrejón *et al*, 2021; Llorens-Bobadilla *et al*, 2015). Quiescent GSCs evade anti-mitotic therapies and hijack NSC-like properties to drive tumour re-growth (Deleyrolle *et al*, 2011; Ishii *et al*, 2016; Chen *et al*, 2012). Thus, understanding the mechanisms controlling GSC quiescence will be important for the design of rational therapeutic strategies that might suppress patient relapse.

NSCs expanded in culture have overlapping gene regulatory networks with GBMs and provide a genetically tractable experimental *in vitro* model that has been useful in delineating the pathways controlling GSC quiescence (Ying *et al*, 2003; Sun *et al*, 2008; Conti *et al*, 2005; Pollard *et al*, 2009; Bulstrode *et al*, 2017; Carén *et al*, 2015; Bressan *et al*, 2021; Marqués-Torrejón *et al*, 2021). Bone-morphogenetic protein 4 (BMP4) induces quiescence of NSCs *in vitro* and *in vivo* (Bond *et al*, 2012; Martynoga *et al*, 2013; Mira *et al*, 2007; Sun *et al*, 2011; Marqués-Torrejón *et al*, 2021), while the mitogens EGF and FGF-2 stimulate proliferation. Previously, we demonstrated that overexpression of the GBM-associated master regulators FOXG1 and SOX2 drives quiescent mouse NSCs into a proliferative radial glia-like state (Bulstrode *et al*, 2017) and induces transcriptional changes at many key cell cycle and epigenetic regulators. In particular, *FoxO3*, which induces quiescence and prevents premature NSC differentiation, is directly repressed by FOXG1 (Renault *et al*, 2009; Bulstrode *et al*, 2017). FOXG1 is therefore an important regulator of quiescence in NSCs and GSCs. Determining the genes and pathways operating downstream of elevated FOXG1 will therefore help our understanding of normal NSC development, adult NSC homeostasis and GBM biology.

The FoxO family are key downstream effectors of PI3K-Akt signalling, controlling genes governing diverse cellular processes including proliferation, metabolism, differentiation, and apoptosis. While FoxO factors can have context-dependent roles in supporting cellular resilience, they are most well-known for their tumour suppressive functions in tissue homeostasis, ageing and cancer (Hornsveld *et al*, 2018; Dansen & Burgering, 2008). FoxO1/3/4 are broadly expressed during development and adulthood and, while discrete roles have been identified, they appear to regulate common target genes *in vitro* with likely significant redundancies (Paik *et al*, 2007).

FoxO6 is the most recently identified FoxO member; it was initially reported to be expressed mainly within the CNS of adult mammals (Hoekman *et al*, 2006), but may also have roles in other tissues such as liver and muscle (Kim *et al*, 2011). Compared to FoxO1/3/4, it has several unique molecular characteristics:

FoxO6 has a low sequence homology (~30%) to other FoxO factors, lacks one of three consensus PKB phosphorylation sites and the presence of a nuclear export signal is debated (Kim *et al*, 2013). Unlike other FoxO members, FoxO6 does not undergo complete nucleo-cytoplasmic shuttling in response to PI3K-Akt-mediated phosphorylation (van der Heide *et al*, 2005; Jacobs *et al*, 2003). These features suggest a distinct cellular function. Indeed, in several cancers FOXO6 is elevated and has oncogenic roles, triggering increased proliferation and progression (Qinyu *et al*, 2013; Rothenberg *et al*, 2015; Wang *et al*, 2017; Lallemand *et al*, 2018).

Here, we demonstrate that FoxO6 is transcriptionally activated downstream of elevated FOXG1 in both mouse NSCs and GSCs, and is necessary for FOXG1-driven exit from quiescence. Following forced expression of FoxO6, we observed a stimulation of macropinocytosis, a cellular process involved in nutrient uptake that requires Pak1-regulated actin cytoskeleton remodelling. Gain- and loss-of-function mechanistic studies demonstrate Pak1 is upregulated by FOXG1 overexpression and downregulated upon FoxO6 loss in proliferative NSCs. Altogether, these data suggest a functional pro-oncogenic role for FoxO6 in the regulatory transitions, such as cell shape and metabolic changes, that must be initiated as cells exit from quiescence into proliferation.

## Results

### FOXG1 transcriptionally activates *FoxO6* in mouse NSCs and GSCs

Previously we found that overexpression of the master regulators FOXG1 and SOX2 supports cell cycle reentry of quiescent mouse NSCs. ChIP-seq and RNA-seq data identified *FoxO6* as a strong candidate FOXG1/SOX2-regulated target gene (Bulstrode *et al*, 2017). Here, we hypothesise that, in contrast to FoxO3, FoxO6 may have unique roles in supporting proliferation. To investigate the effect of elevated FOXG1 on *FoxO6* expression – thereby mimicking the increased levels seen in GBMs – we used clonal adult mouse NSC lines harbouring a Doxycycline (Dox)-inducible *FOXG1-V5* construct, as reported previously (Bulstrode *et al*, 2017).

Elevated FOXG1-V5 was found to significantly increase *FoxO6* expression in proliferating NSCs (Figure 1A). After 24 h, *FoxO6* levels increased by ~17-fold and ~4-fold in two independent clonal NSC lines, termed ‘F6’ and ‘F11-19’, respectively. To circumvent a lack of FoxO6-specific antibodies, an HA tag was inserted using CRISPR/Cas9-mediated homologous recombination at the 3’ end of *FoxO6* in F6 cells (Figure 1B). In agreement with mRNA upregulation, we observed a clear induction of FoxO6 protein in response to elevated FOXG1 (+Dox) (Figures 1C and D). These data indicate that FoxO6 mRNA and protein expression are activated downstream of FOXG1 in normal NSCs.

**Figure 1.**
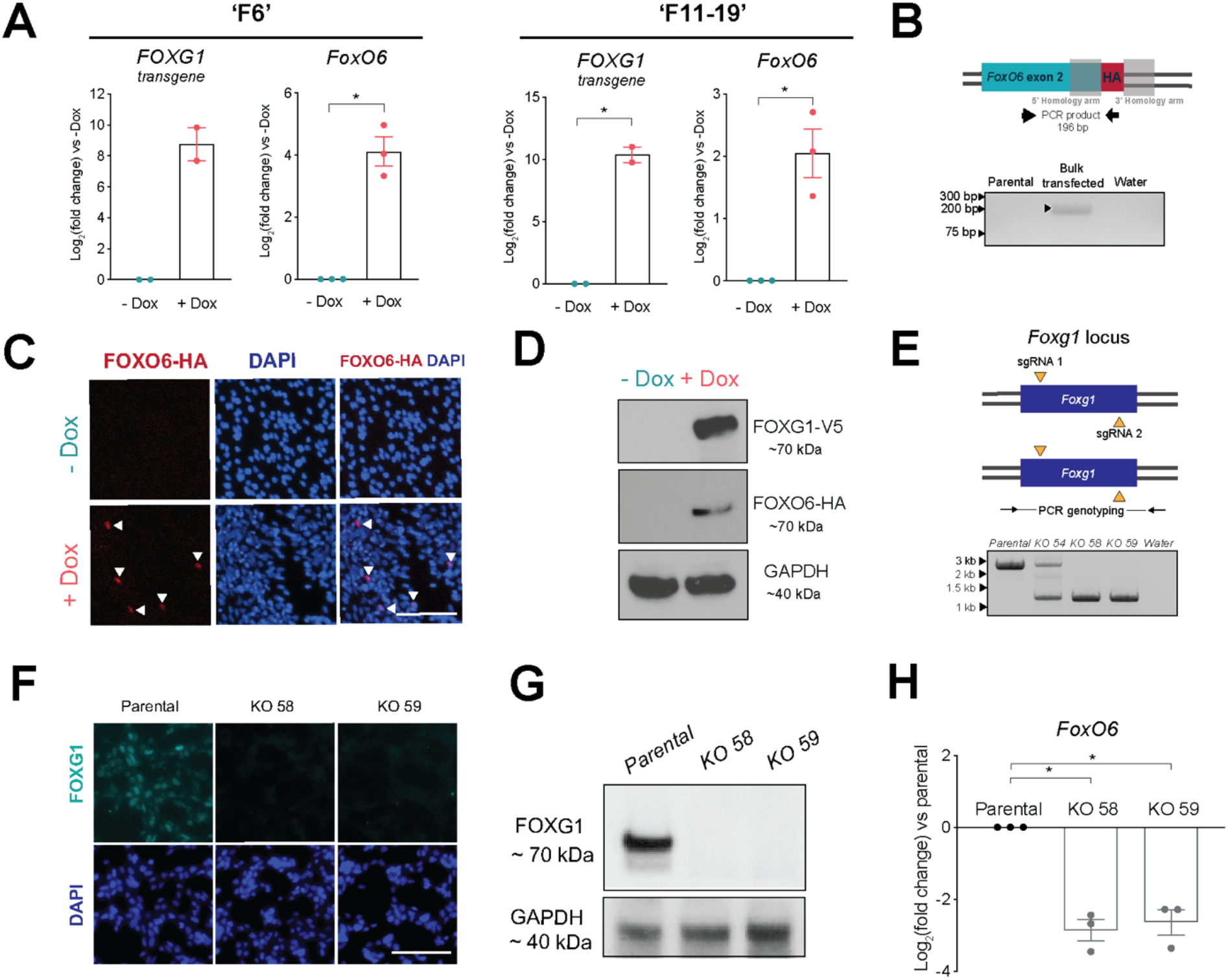
Elevated FOXG1 transcriptionally activates *FoxO6* in mouse NSCs and GSCs. **(A)** qRT-PCR analysis of *FOXG1* transgene and endogenous *FoxO6* expression in two independent adult mouse NSC lines (‘F6’ and ‘F11-19’) with Dox-inducible *FOXG1-V5* expression grown in NSC media with or without Dox for 24 h. Expression shown relative to -Dox (in which log_2_(FC) = 0). Mean +/− SEM. n=2/3 independent experiments, respectively. Each data point shows the mean of one experiment performed in technical duplicates. Two-tailed one sample t-test. * P ≤ 0.05. **(B)** Schematic of HDR-mediated knock-in of an HA epitope tag at the 3’ end of last FoxO6 coding exon in F6 cells. PCR genotyping of the bulk transfected F6 cell population revealed a 196 bp product, indicating the presence of cells with insertion of the HA tag at the 3’ end of FoxO6. **(C)** Wide-field immunofluorescent images following immunocytochemistry (ICC) of FoxO6-HA (red) and DAPI (blue) in the tagged F6 NSCs following Dox addition for 4 days in EGF/FGF-2. Scale bar: 100 μm. **(D)** Western immunoblot analysis of FOXG1-V5 and FoxO6-HA protein expression in tagged F6 NSCs following Dox addition for 4 days in EGF/FGF-2. GAPDH was used as a loading control. **(E)** Experimental strategy for *Foxg1* deletion in IENS-GFP cells. Yellow triangles show the target sites of the sgRNAs at either the 5’ or 3’ end of the coding exon. PCR genotyping of parental IENS-GFP cells and *Foxg1* KO clonal cell lines (KO 58 and KO 59). Wildtype PCR product ~2.6 kb, knockout PCR product ~1.3 kb. **(F)** ICC analysis confirms loss of FOXG1 protein expression in IENS-GFP *Foxg1* KO clonal lines (KO 58 and KO 59). Scale bar: 100 μm. **(G)** Western immunoblot confirming loss of Foxg1 protein in the two independent IENS-GFP knock-out clonal lines (KO 58 and KO 59). GAPDH was used as a loading control. **(H)** qRT-PCR analysis of *FoxO6* expression in IENS-GFP *Foxg1* knock-out clonal lines, compared to parental IENS-GFP (in which log_2_(FC) = 0). Mean +/− SEM. n=3 independent experiments. Each data point shows the mean of one experiment performed in technical duplicates. Two-tailed one sample t-test. * P ≤ 0.05.

We next explored FoxO6 levels in a mouse GBM model cell line (IENS), which expresses FOXG1 at higher levels than mouse NSCs (Bulstrode *et al*, 2017). We tested if *Foxg1* ablation affected *FoxO6* expression. Following CRISPR/Cas9-mediated bi-allelic deletion of *Foxg1* in IENS, a significant decrease (6-7 fold, or 84-86%) in *FoxO6* expression was observed in two independent clonal cell lines (Figures 1E-H). Elevated Foxg1 is therefore necessary for FoxO6 expression in GSCs and is sufficient to induce increased FoxO6 expression in NSCs.

### FOXG1 induction of *FoxO6* occurs early during the exit of NSCs from quiescence

We next assessed *FoxO6* levels during the early phases of NSC exit from quiescence following FOXG1 overexpression. We used quiescence conditions previously shown to induce cell cycle exit, downregulation of NSC markers, and upregulation of quiescent marker expression, namely: treatment with BMP4 at low density for 24 h (Figure 2A) (Bulstrode *et al*, 2017). Following exchange of BMP4 for culture media with EGF and FGF-2, FOXG1 induction (+Dox) drives cells to re-enter cell cycle and form NSC-like colonies (Figure 2B-F). As expected, we find that Dox addition induced a 235-fold upregulation in *FOXG1* expression by 24h compared to the non-BMP treated control (EGF/FGF-2) (Figure 2G). *FoxO6* was dramatically upregulated at these early timepoints, prior to any visible proliferative response, with a ~6.5-fold upregulation by 24h compared to the non-BMP4 treated control, ~16-fold higher than without Dox (Figure 2H). Increased FoxO6 expression is therefore an early part of the response to FOXG1 in the transition from quiescence to proliferation, consistent with it being an important functional downstream effector.

**Figure 2.**
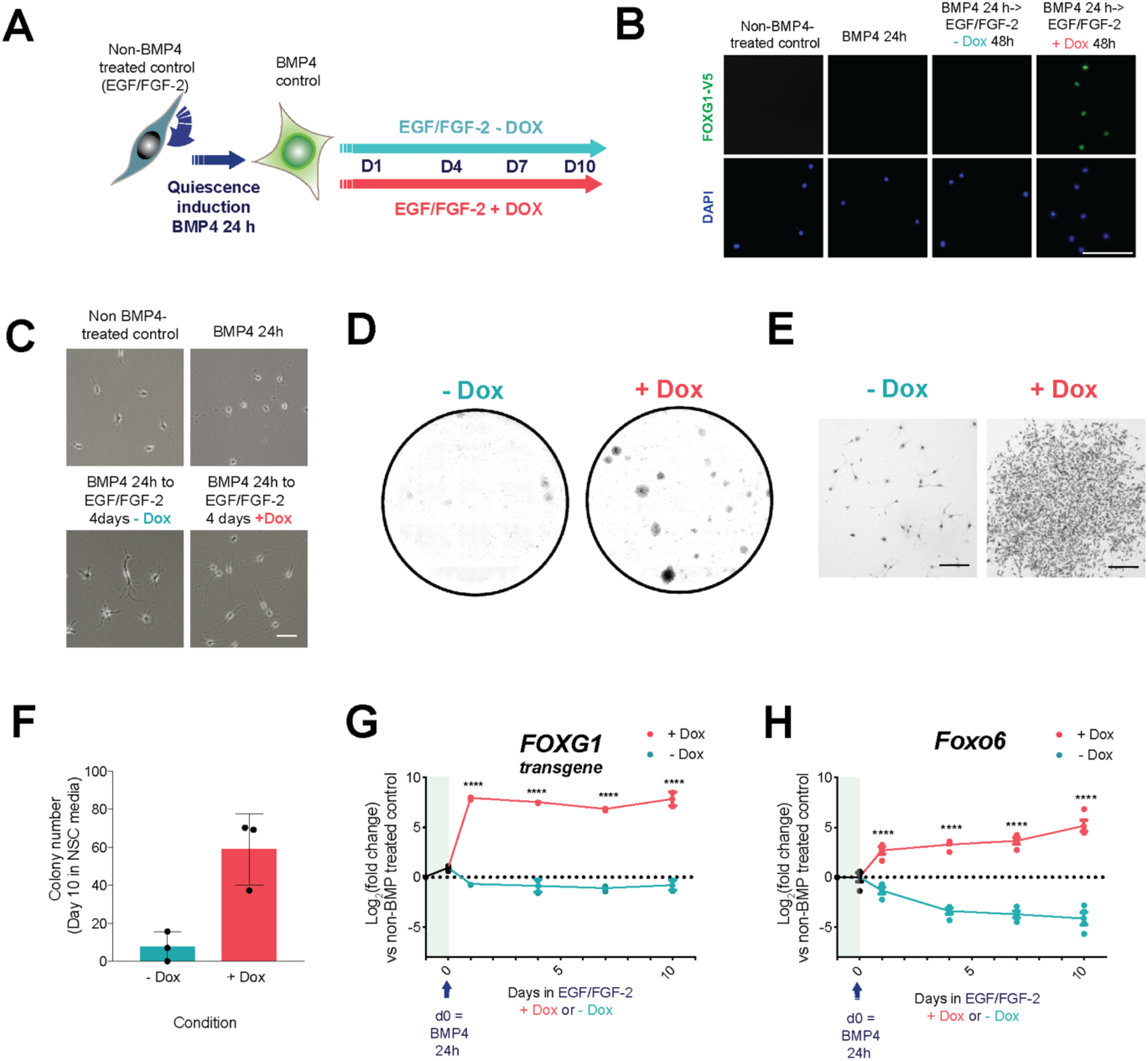
FOXG1 induces *FoxO6* during quiescent NSC reactivation. **(A)** Schematic of the experimental design for assessing FOXG1-induced reactivation of quiescent NSCs and associated changes in gene expression, using clonal ‘F6’ adult mouse NSC line with Dox-inducible *FOXG1-V5* expression. Non-BMP4 treated control = cells in NSC media with EGF/FGF-2. **(B)** ICC for V5, confirming FOXG1-V5 expression upon Dox addition (Scale bar: 100 μm). **(C)** Representative phase-contrast images showing changes in cell morphology upon addition of Dox (Scale bar: 100 μm). **(D)** Colony formation after 24 h BMP4 treatment followed by 10 days in EGF/FGF-2 with or without Dox. Representative images shown of wells stained with methylene blue and imaged on a bright-field microscope. n=3 independent experiments. **(E)** Higher magnification phase-contrast images of representative colonies after 24 h BMP4 treatment and 10 days in EGF/FGF-2 with or without Dox as in panel (D) (Scale bar: 200 μm). **(F)** Number of colonies formed after 24 h BMP4 treatment and 10 days in EGF/FGF-2 with or without Dox. Mean +/− SD, n=3 independent experiments. Each data point shows the mean of one experiment performed in technical triplicates. **(G)** qRT-PCR analysis of human *FOXG1* transgene and **(H)** *FoxO6* expression during the reactivation time course. Pink = + Dox addition, Blue = No Dox addition. Expression shown relative to non BMP4-treated (EGF/FGF-2) control (in which log_2_(FC)= 0 (dotted line)). Day 0 = expression after 24 h BMP4 treatment. Mean +/− SEM. n = 2 (*FOXG1*) or 4 (*FoxO6*). Each data point shows the mean of one experiment performed in technical duplicates. **** P≤0.0001. Two way Anova with Sidak correction.

### FOXG1-induced reactivation of quiescent NSCs is significantly impaired upon *FoxO6* loss

To assess whether *FoxO6* was required for FOXG1-induced quiescence exit, CRISPR/Cas9 was used to generate clonal adult mouse NSC lines with bi-allelic deletion of the first *FoxO6* coding exon (Figure S1A-B). qRT-PCR analysis confirmed loss of *FoxO6*, with a >98% decrease in expression following CRISPR treatment compared to untargeted parental cells (Figure 3A).

**Figure 3.**
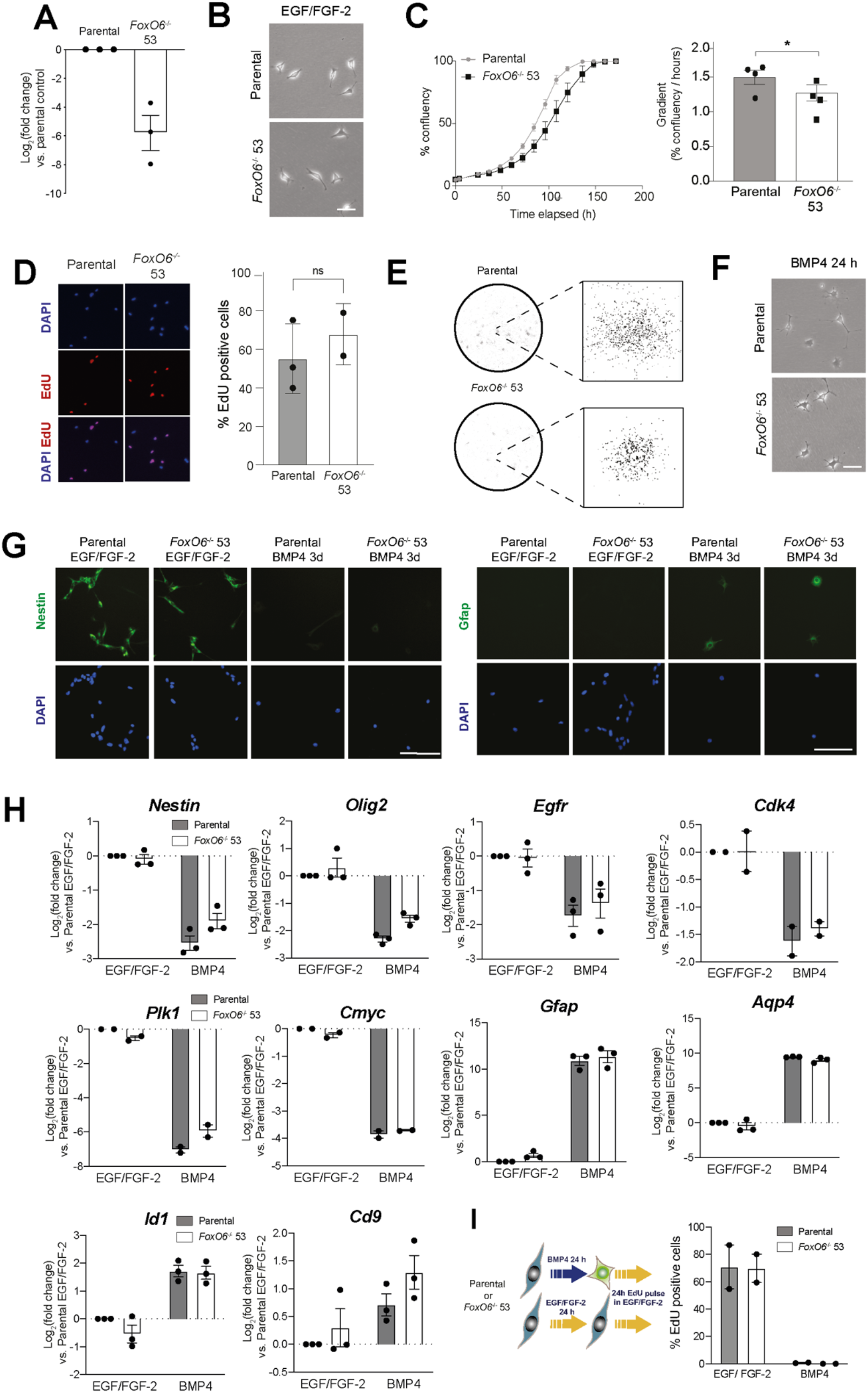
FoxO6 is not required for continued NSC proliferation or response to BMP4. **(A)** qRT-PCR analysis of *FoxO6* mRNA levels in *FoxO6* KO clonal cell line 53, compared to ANS4 parental cells (in which log_2_(FC) = 0). Expression values were normalised to Gapdh. Y axis represents log_2_(Fold change). Mean +/− SEM. n=3 independent experiments. Each data point shows the mean of one experiment, performed in technical duplicates. **(B)** Representative phase-contrast images showing typical NSC morphology in parental and *FoxO6* KO 53 cells. Scale bar: 25 μm. **(C)** Growth curve analysis of parental and *FoxO6* KO 53 clonal cells in EGF/FGF-2. Light grey = parental, black = FoxO6 KO 53. Mean +/− SD, n=3 technical replicates. Representative of n=4 independent experiments. Graph showing the gradient of the linear portion of the logistic growth curve (% / h). Mean +/− SEM, n=4 independent experiments. Two-tailed paired Student’s t-test. * P ≤ 0.05. **(D)** EdU incorporation assay (24h pulse) in EGF/FGF-2 for parental and FoxO6 KO 53 cells. (Left) Representative fluorescent images of EdU incorporation after 24 h pulse. (Right) Plot shows mean +/− SEM, n=3 independent experiments. Each data point shows the mean of one experiment performed in technical triplicates. **(E)** Brightfield images of colony formation by parental or FoxO6 KO 53 cells 10 days after plating at low density in NSC media (EGF/FGF-2). Plates stained with methylene blue. **(F)** Representative phase-contrast images showing morphology of ANS4 and FoxO6 KO 53 after 24h BMP4 treatment. Scale bar: 25 μm. **(G)** ICC analysis of Nestin and Gfap expression in ANS4 parental and FoxO6 KO 53 cells in EGF/FGF-2 or 3 days BMP4 treatment at low density. Scale bar: 100 μm. **(H)** qRT-PCR analysis of NSC (*Nestin, Olig2, Egfr*), cell cycle (*Plk1, Cdk4, Cmyc*), and astrocyte/quiescence (*Gfap, Aqp4, Id1, Cd9*) markers, in parental and FoxO6 KO 53 cells in EGF/FGF-2 and after 24 h BMP4 treatment. Expression shown relative to parental in EGF/FGF-2 (in which log_2_(FC) = 0). Mean +/− SEM. n=2/3 independent experiments. Each data point shows the mean of one experiment performed in technical duplicates. **(I)** EdU incorporation after treatment with BMP4 or EGF/FGF-2 for 24h, followed by a 24h EdU pulse in EGF/FGF-2. Mean +/− SEM. n=2 independent experiments. Each data point shows the mean one experiment, performed in technical triplicates.

*FoxO6*^-/-^ clonal cells displayed a bipolar phase-bright morphology characteristic of proliferative NSCs (Figure 3B). Although confluence analysis suggested a marginally reduced growth compared to parental cells, EdU incorporation did not indicate any significant changes in proliferation (Figure 3C-D). Furthermore, *FoxO6^-/-^* cells were found to form typical NSC colonies in EGF/FGF-2 (Figure 3E). This indicated that FoxO6 is not necessary for NSC proliferation or colony formation under optimal self-renewing conditions.

Following BMP4 treatment at low density for 24hr, both parental and *FoxO6*^-/-^ NSCs displayed a characteristic change to a stellate astrocytic morphology (Figure 3F). ICC analysis confirmed a decrease in NESTIN and increase in GFAP expression in both parental and *FoxO6^-/-^* NSCs after 3 days of BMP4 treatment (Figure 3G). qRT-PCR analyses showed upregulation of astrocytic/quiescence markers (*Gfap, Aqp4, Id1, Cd9*) and downregulation of NSC and cell cycle markers (*Nestin, Olig2, Egfr, cMyc, Plk1, Cdk4*) in parental and *FoxO6^-/-^* cells (Figure 3H). Cell cycle exit following BMP4 treatment in both parental and *FoxO6^-/-^* cells was confirmed by loss of EdU incorporation (Figure 3I). Proliferation and quiescence entry analyses in additional *FoxO6^-/-^* clonal lines showed consistent results, with only one out of three KO lines displaying altered proliferation (Figure S1C-E). These results suggest that NSC identity is not lost following *FoxO6* disruption, and that *FoxO6* is not required for cytostatic BMP response and entry into the quiescent state.

We next tested whether FoxO6 was essential for exit from quiescence. Parental and *FoxO6^-/-^* NSCs were transfected with the Dox-inducible FOXG1-V5 overexpression construct using the PiggyBac transposase system. Quiescence exit was assessed following BMP4 treatment and return of cells to EGF/FGF-2 for 10 days, with or without FOXG1-V5 induction (-/+ Dox), as previously mentioned. Similar levels of transgene induction were achieved in both parental and *FoxO6*^-/-^ populations with inducible FOXG1, as assessed by FOXG1-V5 qRT-PCR and ICC (Figure 4A-C). Strikingly, *FoxO6^-/-^* cells showed dramatically reduced capacity to reform proliferative NSC colonies (Figure 4D-F) and, unlike the parental cells, did not highly upregulate the NSC marker Nestin or proliferative marker Ki67 upon FOXG1 induction (Figure S2A). *FoxO6^-/-^* NSCs showed striking morphological changes from the typical ‘fried-egg’, stellate astrocytic morphology, to an elongated spindle shape that was distinct from the typical bipolar phase-bright morphology of parental cells with FOXG1 induction (Figure S2B). After an extended period (25 days) only minor evidence of colony formation was seen in the *FoxO6^-/-^* population (Figure S2C). Together these data suggest *FoxO6* is a key downstream effector of elevated FOXG1, required for efficient transition from quiescence to proliferation. Without FoxO6, quiescent cells fail to undergo the shape changes and cell cycle re-entry typical of quiescence exit in NSCs.

**Figure 4.**
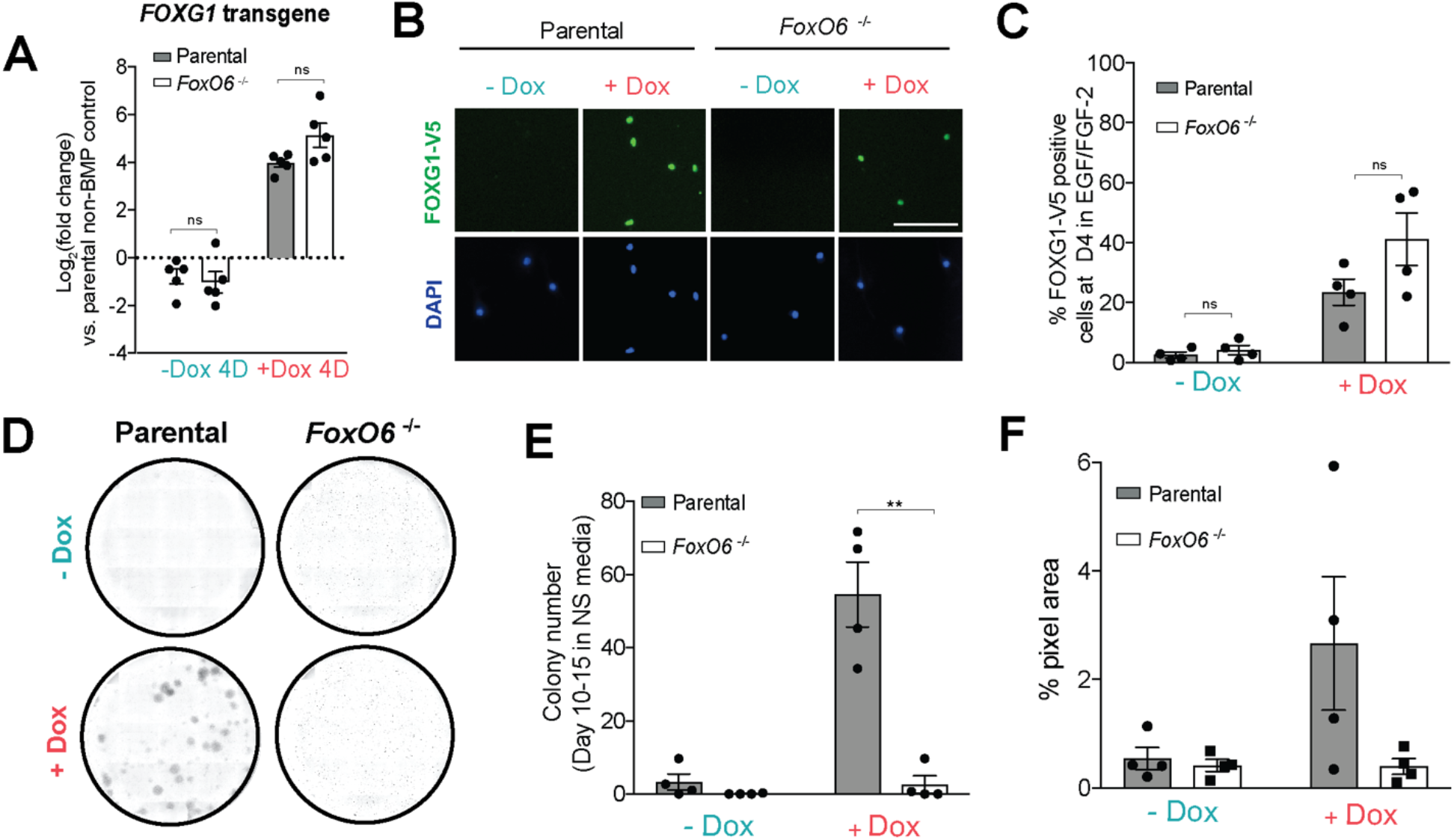
FOXG1-induced reactivation of quiescent NSCs is inhibited upon *FoxO6* loss. **(A)** qRT-PCR analysis of *FOXG1* transgene expression in parental or FoxO6 KO 53 cells engineered with inducible FOXG1-V5 after 24 h BMP4 and return to EGF/FGF-2 with our without Dox for 4 days. Expression shown relative parental non-BMP treated (EGF/FGF-2) control (in which log_2_(FC) = 0 (dotted line)). Mean +/− SEM. n=5 independent experiments. Each data point shows the mean of one experiment performed in technical duplicates. Two-tailed paired t-test. **(B)** Representative ICC images showing FOXG1-V5 expression after 24 h BMP4 and 4 days in EGF/FGF-2, with or without Dox, in parental and FoxO6 KO 53 cells with inducible FOXG1-V5. Scale bar: 100 μm. **(C)** Percentage of parental or FoxO6 KO 53 cells with inducible FOXG1 expressing FOXG1-V5 (assessed by ICC) after 24 h BMP4 and 4 days in EGF/FGF-2, with or without Dox. Mean +/− SEM. n = 4 independent experiments. Each data point shows the mean of one experiment performed in technical triplicates. Twotailed paired t-test. **(D)** Representative images of colony formation assay with parental and FoxO6 KO 53 cells at day 10 in EGF/FGF-2, with or without Dox. Plates stained with methylene blue following fixation. **(E)** Numbers of colonies formed after 24 h BMP4 and 10-15 days in EGF/FGF-2, with or without Dox as in panel (D). Mean +/− SEM, n= 4 independent experiments. Each data point shows the mean of one experiment performed in technical triplicates. Two tailed paired Student’s t-tests. ** P ≤ 0.01. **(F)** Percentage of the well area covered by cells after 24 h BMP4 and 10-15 days in EGF/FGF-2, with or without Dox as in panel (D). Mean +/− SEM. n=4 independent experiments. Each data point shows the mean of one experiment performed in technical triplicates.

### Elevated FoxO6 induces the formation of large acidic vacuoles by macropinocytosis

To explore the specific pathways through which FoxO6 might operate to stimulate quiescence exit, we next tested the effects of forced FoxO6 expression. We established adult mouse NSCs with Dox-inducible *FoxO6-HA-IRES-mCherry* overexpression using the PiggyBac transposase system. Following FACS enrichment of mCherry positive cells, FoxO6-HA overexpression was confirmed by Western blotting, ICC and qRT-PCR analysis (Figure 5A-B).

**Figure 5.**
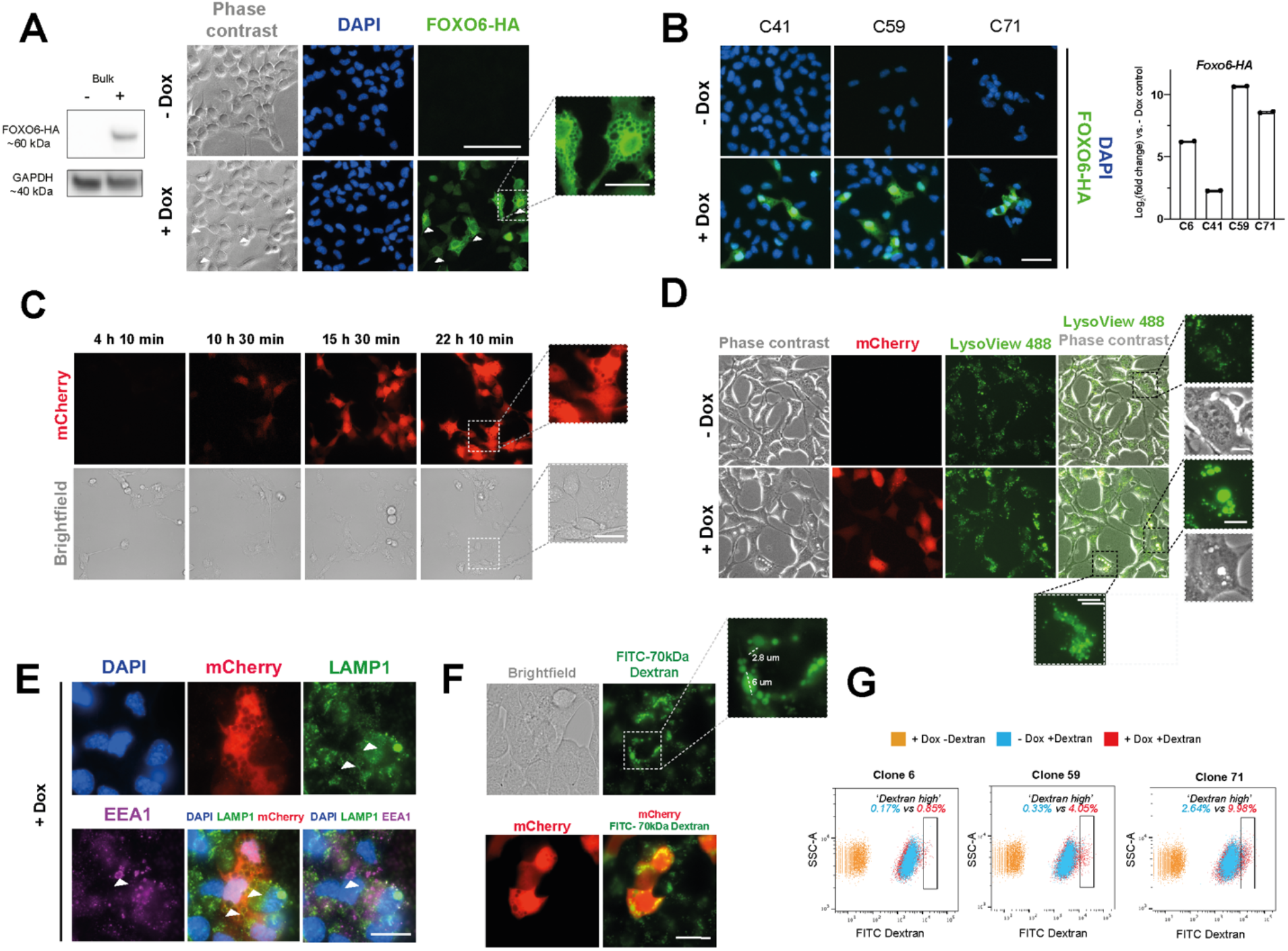
Elevated FoxO6 induces the formation of large acidic vacuoles by macropinocytosis. **(A)** Western blotting and ICC analysis confirming expression of FoxO6-HA transgene following 24h Dox treatment. Image inset highlights appearance of vacuoles in FoxO6-HA-overexpressing cells (+Dox). Scale bars are 100 um or 25um. GAPDH is used as a housekeeping loading control. **(B)** ICC of FoxO6-HA overexpression showing vacuolisation upon Dox addition to clonal NSC lines with Dox-inducible *FoxO6-HA-IRES-mCherry* expression. ICC scale bar = 50 uM. qRT-PCR for FoxO6-HA expression in clonal cell lines (Mean +/− SD, technical duplicates, -Dox = 0 for each clonal line). **(C)** Live imaging following Dox addition to clonal NSCs with Dox-inducible *FoxO6-HA-IRES-mCherry* expression (C71). Dox was added 4 hours prior to imaging. Images were obtained every 10 minutes for ~18 hours. Scale bar 100 um or 25 um. **(D)** Imaging of LysoView-488 accumulation in clonal FoxO6-inducible cell line (C71) with or without Dox addition (2 days). Scale bars 50 um and 10 um. **(E)** ICC for lysosomal marker LAMP1 and early endosomal marker EEA1 to visualise co-localisation with vacuoles (C59, following overnight Dox incubation). Scale bar 25 um. **(F)** Live imaging of 70 kDa FITC-dextran uptake and mCherry expression following incubation with Dox overnight in FoxO6-inducible cell line (C59). Scale bar 25 um. **(G)** Flow cytometry-based quantification of 70 kDa FITC-dextran uptake following incubation with Dox overnight in FoxO6-inducible cell lines (6, 59, 71). Samples displayed are +Dox -Dextran control, -Dox + Dextran and +Dox +Dextran. Gating shows ‘Dextran high’ population. Percentages represent the increase in ‘Dextran high’ cells upon Dox addition.

We used these transfected cells to explore cellular responses to elevated FoxO6 by microscopy. Strikingly, prominent vacuolisation was observed upon FoxO6 overexpression across multiple clonal lines (Figure 5A-B). Live cell imaging across a time-course revealed vacuole formation occurred within 10-11 hours of Dox addition, coincident with mCherry expression (Figure 5D). Neither treatment of untransfected adult mouse NSCs with Dox nor overexpression of an alternative transcription factor (using the same plasmid constructs with only the gene of interest substituted) resulted in vacuole formation, indicating that the phenotype was specific to FoxO6 overexpression and not a result of Dox treatment, the HA tag or mCherry overexpression (Figure S3A-B).

We next characterised the resulting vacuoles using various imaging methods. We first ruled out these structures being lipid droplets or enlarged lysosomes, which have been implicated in quiescence regulation (Ramosaj *et al*, 2021; Leeman *et al*, 2018). Staining with the neutral lipid dye BODIPY did not reveal colocalisation with the vacuoles (Figure S3C). In contrast, following LysoView™ incubation, we observed strong fluorescent signal co-localised with the vacuoles indicating their acidification (Figure 5D). Interestingly, not all structures showed equal LysoView™ accumulation, suggesting they were at different stages of acidification and maturation, and therefore not simply enlarged lysosomes. Consistently, ICC for the lysosomal membrane marker, LAMP1, and the early endosomal marker, EEA1, did not reveal uniform co-localisation with the vacuoles (Figure 5E) and Western blot analysis showed no clear increase in EEA1 nor LAMP1 expression upon FoxO6-HA induction (Figure S3E).

Uptake of a fluorescent EGF ligand revealed much smaller puncta, indicating the vacuoles were not formed by receptor-mediated endocytosis (Figure S3D). Instead, the vacuole size (as large as 6 microns in diameter (Figure 5F)) was strongly suggestive of non-selective macropinocytosis – an actin-driven process by which extracellular contents are engulfed and processed along the endosomal pathway (Swanson & Watts, 1995; Lim & Gleeson, 2011; Kerr & Teasdale, 2009). To test this hypothesis, we incubated cells with large molecular weight 70kDa FITC-dextran, a well-established marker of macropinocytosis (Galenkamp *et al*, 2019; Commisso *et al*, 2014). In cells treated overnight with Dox, clear FITC-dextran uptake was visible within the vacuoles (Figure 5F). Flow cytometry quantification of FITC-dextran uptake confirmed an increase in the percentage of ‘Dextran high’ cells following Dox addition compared to -Dox controls across three FoxO6-HA inducible cell lines (Figure 5G and S3F).

EdU analysis suggested vacuolisation did not per se provide a growth advantage; instead, highly vacuolated cells were associated with EdU negativity following a 24h pulse (Figure S3G-H). The vacuolated cells could be passaged and remained in culture (Figure S3I), ruling out a novel form of cell death induced by hyperactivated macropinocytosis in cancer named methuosis (Song *et al*, 2021; Overmeyer *et al*, 2008). Together these observations suggest macropinocytosis, or the pathways that stimulate it, may be an important feature of FoxO6 activity as cells exit quiescence. Macropinocytosis in cancer is associated with nutrient acquisition to aid proliferation (Recouvreux & Commisso, 2017; Commisso *et al*, 2013). The lack of proliferative advantage conferred by FoxO6 overexpression is consistent with the need for other supporting pathways downstream of FOXG1 for quiescence exit.

### Pak1 expression is upregulated upon FOXG1 elevation and downregulated upon FoxO6 loss in proliferative NSCs

The p21 (Cdc42/Rac)-activated kinase, Pak1, is a specific regulator of macropinocytosis controlling actin cytoskeleton dynamics (Dharmawardhane *et al*, 2000), and has been reported as a FoxO6 target in the transcriptional pathway orchestrating neuronal polarity (De La Torre-Ubieta *et al*, 2010). Both FoxO6 and Pak1 have published roles in memory consolidation and synaptic function (Salih *et al*, 2012; Civiero & Greggio, 2018).

To investigate a potential involvement of Pak1, we explored if its levels were modulated by FOXG1 or FoxO6. qRT-PCR and Western blot analysis revealed an increase in Pak1 expression in proliferating NSCs (compared to the -Dox control) upon FOXG1 induction (Figures 6A-B). Furthermore, Pak1 expression was decreased in *FoxO6^-/-^* proliferative NSCs by both qRT-PCR and Western blot analysis (Figure 6C-E). This suggests that in proliferative culture conditions, FoxO6 is needed to sustain Pak1 expression, consistent with a potentially important role in the earliest phases of cell cycle re-entry during quiescence exit. Finally, qRT-PCR analysis of *FOXG1, FoxO6* and *Pak1* following FOXG1-induced quiescence exit (in F6 cells) revealed higher levels of *Pak1* in +Dox compared to the -Dox control, coincident with *FOXG1* inducing *FoxO6* upregulation (Figure 6F). In summary, our findings show that FOXG1-mediated induction of FoxO6 is required for efficient quiescence exit of NSCs. We speculate this might operate through modulation of a Pak1-macropinocytosis-related pathway that is initiated as cell undergo regulatory changes (such as shape and nutrient requirements) in preparation for cell cycle re-entry.

**Figure 6.**
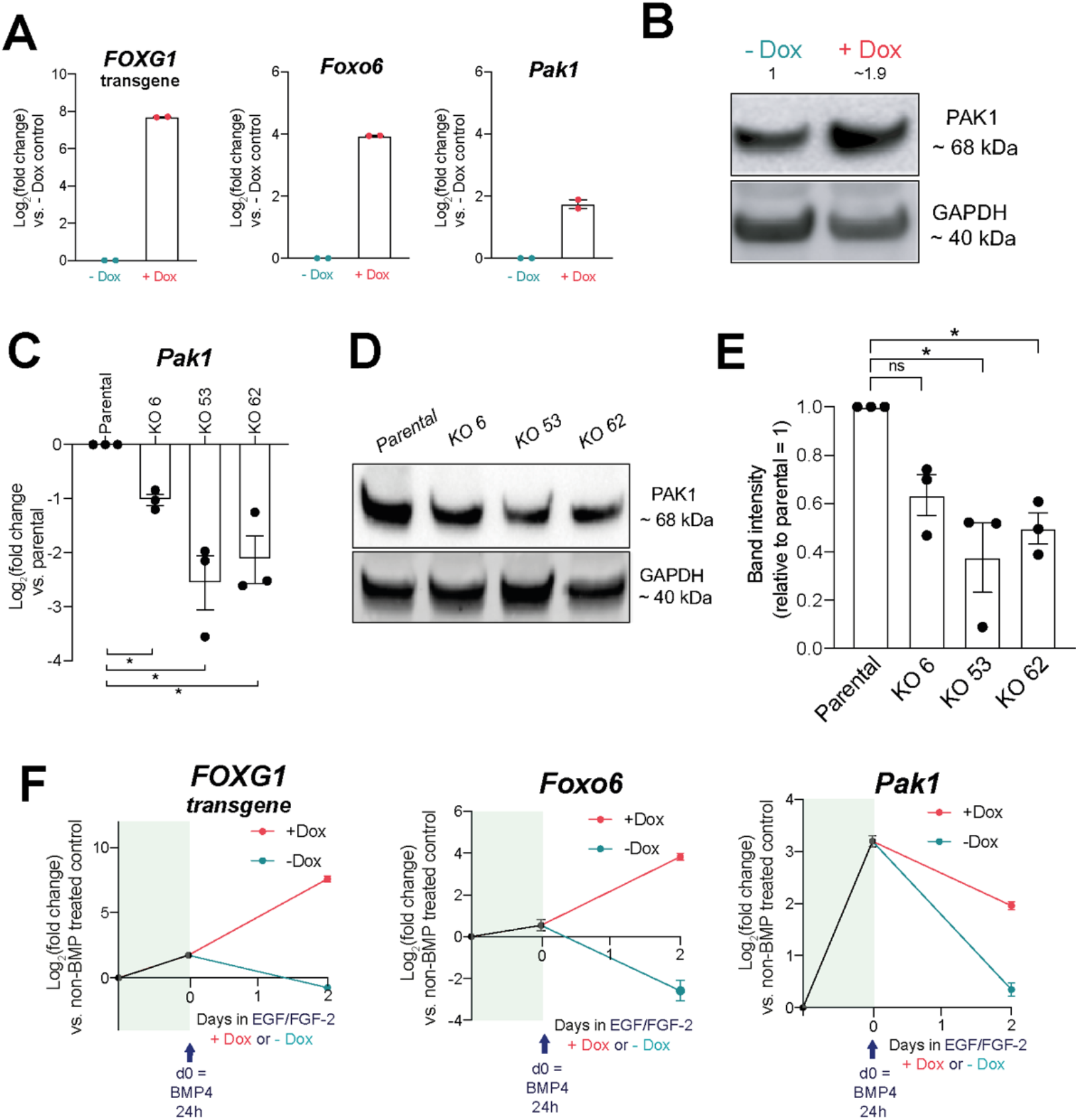
Pak1 expression is upregulated upon FOXG1 elevation and downregulated upon FoxO6 loss in proliferative NSCs. **(A)** qRT-PCR analysis of *FOXG1* transgene, and endogenous *FoxO6* and *Pak1* expression in F6 cells with Dox-inducible FOXG1-V5 grown in EGF/FGF-2 for 24h plus or minus Dox (n=2 biological replicates, Mean +/− SEM. Each data point shows the mean of one experiment performed in technical duplicates). **(B)** Western blot analysis of Pak1 expression in F6 cells treated with Dox in EGF/FGF for 24h. GAPDH is used as a loading control. Quantification of Pak1 bands normalised to GAPDH and -Dox control shown, where -Dox =1. **(C)** qRT-PCR analysis of *Pak1* expression in ANS4 parental versus FoxO6 KO clones 6, 53 and 62 (n=3 biological replicates, Mean +/− SEM. Each data point shows the mean of one experiment performed in technical duplicates.) Two-tailed one sample t-test, * p<0.05. **(D)** Western blot analysis of Pak1 expression in parental versus FoxO6 KO clones 6, 53 and 62 (n=3 biological replicates). **(E)** Quantification of Pak1 Western blot band intensities as in panel (D). Parental = 1 as shown by the dotted line (n=3 biological replicates, Mean +/− SEM. Each data point shows the intensity from one experiment). Two-tailed one sample t-test, * p<0.05. **(F)** qRT-PCR analysis of *FOXG1* transgene, and endogenous *FoxO6* and *Pak1* expression in F6 cells after 24 h BMP4 and return to EGF/FGF-2 with or without Dox for 2 days. Expression shown relative non BMP-treated (EGF/FGF-2) control (in which log_2_(FC) = 0). Day 0 = expression after 24 h BMP4 treatment (n=2 biological replicates, Mean +/− SEM. Each data point shows the mean of one experiment performed in technical duplicates).

## Discussion

Understanding the molecular mechanisms governing control of NSC quiescence has important implications in GBM, a highly aggressive adult brain cancer in which quiescent NSC-like stem cells drive relapse. Our findings here extend our previous observations that high levels of FOXG1 and SOX2 drive a proliferative radial glial-like NSC phenotype, in part through repression of the tumour suppressor *FoxO3* (Bulstrode *et al*, 2017). Here, we show that *FoxO6*, an underexplored FoxO member, is a downstream target activated by elevated FOXG1.

Whilst FoxO3 has been well-described as a tumour suppressor that preserves NSC quiescence (Renault *et al*, 2009; Liu *et al*, 2018), our data suggest FoxO6 has an opposite, pro-proliferative, role in FOXG1-induced quiescence exit of NSCs. *FoxO6* loss did not impair NSC proliferation or BMP4-induced quiescence entry. We observed NSC and cell cycle marker downregulation, astrocytic/quiescence marker upregulation, morphological changes and cell cycle exit upon BMP4 treatment, all indicating entry into a quiescent state (Codega *et al*, 2014; Dulken *et al*, 2017; Llorens-Bobadilla *et al*, 2015). However, FoxO6 loss was found to significantly impair FOXG1-induced exit from quiescence. Elevated FOXO6 has indeed been associated with stimulating proliferation and progression in several cancers (Qinyu *et al*, 2013; Rothenberg *et al*, 2015; Wang *et al*, 2017; Lallemand *et al*, 2018). FoxO6 has also been reported to transcriptionally control SOX2, STAT3 and Hippo signalling, all reported to control NSC and GSC self-renewal or proliferation (Ganguly *et al*, 2018; Yang *et al*, 2016; Salih *et al*, 2012; Rothenberg *et al*, 2015; Bulstrode *et al*, 2017; Sun *et al*, 2018).

Our functional studies of FoxO6 suggested that forced expression alone will trigger macropinocytosis – a process involving Pak1-regulated actin cytoskeleton remodelling. Together with literature on both FoxO6 and Pak1 in neuronal polarity and synaptic function (De La Torre-Ubieta *et al*, 2010; Salih *et al*, 2012; Civiero & Greggio, 2018), this led us to investigate Pak1 levels in relation to FOXG1 and FoxO6. Our data suggest in proliferating NSCs (with mitogens EGF and FGF-2), FoxO6 is required to sustain Pak1 expression and FOXG1 induction can result in even higher Pak1 levels. Whilst this is not functionally important in sustaining NSC proliferation, as shown by the lack of proliferation defects upon Foxg1 or FoxO6 deletion (Bulstrode *et al*, 2017), these changes in Pak1 levels may impact regulatory transitions, such as cell shape and metabolic requirements, as cells prepare to exit quiescence into the proliferative radial-glia like state. Our data lead us to speculate a working model in which FoxO6 is activated downstream of FOXG1, and in turn will trigger a Pak1-related pathway that alters actin dynamics and related cell shape/nutrient sensing pathways to facilitate exit from quiescence.

As vacuolisation was not observed upon FOXG1 overexpression (Figure S3J), it is possible that FoxO6-induced macropinocytosis represents a stalled state, with other pathways downstream of FOXG1 necessary to be activated concomitantly to ensure cell cycle re-entry, e.g., through increased pinocytic flux that cannot be assessed within our experimental timeframes. Indeed, active Pak1 has been found to modulate pinocytic cycling, enhancing both FITC-dextran uptake and efflux (Dharmawardhane *et al*, 2000). It is plausible that such an enhancement in pinocytic cycling may aid rewiring of the metabolome required for the transition from quiescence to proliferation (Lee *et al*, 2017; Wani *et al*, 2022; Adusumilli *et al*, 2021). This will require further deeper exploration in future studies. Alternatively, hyperactivation of signalling pathways upon FoxO6 overexpression may result in macropinocytosis as a metabolic stress response. Hyperactivation of Ras signalling, canonical Wnt and PI3K signalling have all been shown to play roles in inducing macropinocytosis (Overmeyer *et al*, 2008; Tejeda-Muñoz *et al*, 2019; Recouvreux & Commisso, 2017). Interestingly, FOXG1 was recently found to synergise with Wnt signalling in driving quiescence exit in GBM (Robertson *et al*, submitted). The activity of FoxO factors is controlled by phosphorylation downstream of IGF/PI3K/AKT signalling (Hay, 2011; Jiramongkol & Lam, 2020). Pak1 is upregulated in various cancer types (Huynh et al, 2015), integrates various signalling pathways, such as PI3K and Ras, and has been reported to phosphorylate and inactivate FoxO1 in breast cancer (Mazumdar *et al*, 2003) and Foxo6 in liver ageing (Kim *et al*, 2015). It is therefore also possible that FoxO6 elevation results in signalling activation that in turn reinforces phosphorylation and deactivation of FoxO tumour suppressors.

FoxO factors are known to modulate metabolic functions in homeostasis and cancer (Yadav *et al*, 2018; Kim *et al*, 2011; Chung *et al*, 2013; Paik *et al*, 2007). FoxO3 protects NSCs against oxidative stress and controls their glucose metabolism to ensure optimal self-renewal (Yeo *et al*, 2013; Renault *et al*, 2009), in part through *Myc* inhibition (Peck *et al*, 2013). In contrast, FoxO6 promotes gastric cancer cell proliferation through *c-Myc* induction (Qinyu *et al*, 2013), and its loss inhibits colorectal cancer cell proliferation, invasion and glycolysis, with decreased PI3K/Akt/mTOR pathway activation (Li *et al*, 2019). In GBM, mTORC2 signalling controls glycolytic metabolism through inhibition of FoxO1/3 and de-repression of *c-Myc* (Masui *et al*, 2013). Elevated FOXG1, itself implicated in regulating mitochondrial functions (Pancrazi *et al*, 2015), may therefore alter FoxO3 and FoxO6 expression to result in deregulated energetics that drive a proliferative state and/or oppose quiescence. Macropinocytosis in cancer has been reported to aid nutrient uptake (Recouvreux & Commisso, 2017; Commisso *et al*, 2013); the role of FoxO6 in linking GSC state transitions with metabolism will therefore be an interesting avenue for further exploration.

With respect to normal NSCs, FoxO6’s roles in the developing and adult brain are less well-defined than FoxO3 (Sun *et al*, 2018; Salih *et al*, 2012). The changing spatial pattern of FoxO6 expression during mouse brain development suggests different functions at distinct stages; yet, the NSC number at E18 is unchanged in *FoxO6* null mice (Hoekman *et al*, 2006; Paap *et al*, 2016). Cortical FoxO6 levels decrease significantly after birth, with adulthood expression regulating synapse formation in the hippocampal CA1/3 regions, as well as cerebellar neuronal polarity (Salih *et al*, 2012; De La Torre-Ubieta *et al*, 2010). Like Foxg1, FoxO6’s homeostatic roles may therefore be subtle in adulthood, and mostly involved in neural plasticity (Yu *et al*, 2019). This is in keeping with our finding that basal FoxO6 levels are low in adult NSCs and not required for sustained proliferation, but are important for cell state transitions. If the FoxO6 levels activated by elevated FOXG1 represent an acquired dependency of GBM, there may be a therapeutic window to target this pathway. However, given the poorly understood roles of FoxO6, further work is needed to determine its specific value as a therapeutic target. Regardless, the balance between these three FOX family members – Foxg1, FoxO3 and FoxO6 – has been revealed by our studies, and others, to be a key signalling node in the context of GBM quiescence control and warrants further investigation.

## Author contributions

K.M.F. conducted experiments, data analysis, prepared the figures and co-wrote the manuscript. C.B. performed western blotting and aided Pak1 investigation. C.G.D. derived the FoxO6 KO cell lines. H.B. derived F6 cell line and preliminary data on FoxO6. R.B.B provided plasmid for FOXG1 KO in IENS, strategy for CRISPR FoxO6 deletion with C.G.D and manuscript editing. K.M. generated plasmid for FoxO6-HA-IRES-mCherry overexpression, and provided NPE-MYEF2-IRES-mCherry control cells. S.M.P. conceived, designed, and coordinated the study and co-wrote the manuscript.

## Acknowledgements

K.M.F. was supported by a studentship from Cancer Research UK (A19680). S.M.P was supported by a Cancer Research UK Senior Research Fellowship (A17368). H.B. was supported by a Wellcome Trust Clinician Research Training Fellowship. R.B.B. was supported by a PhD fellowship from the Science Without Borders Program (CAPES, Governo Dilma Rousseff, Brazil). We acknowledge the Dr Matthieu Vermeren and the Imaging Facility, and Dr Fiona Rossi and the Flow Cytometry Facility at the Centre for Regenerative, University of Edinburgh, for their technical support. We thank Dr Noor Gammoh and Dr Sonja Vermeren for their advice on macropinocytosis and for providing reagents. K.M.F. thanks Dr Maria Angeles Marques-Torrejon, Dr Ester Gangoso and Dr Pooran Singh Dewari for supervision and advice on methodologies, and Vivien Grant for technical assistance. We thank Dr Maria Angeles Marques-Torrejon and Dr Faye Robertson for critical reading of the manuscript. For the purpose of open access, the author has applied a Creative Commons Attribution (CC BY) licence to any Author Accepted Manuscript version arising from this submission.

## Experimental Procedures

### Cell culture

Mouse NSC lines were derived from adult SVZ as described previously (Conti *et al*, 2005; Sun *et al*, 2008). IENS cells, described previously with Ink4a/ARF deletion and EGFRvIII overexpression (Bruggeman *et al*, 2007; Bulstrode *et al*, 2017), were kindly provided by Prof M. Van Lohuizen (NKA, Amsterdam). Established lines were cultured in an adherent monolayer on uncoated tissue culture plastics, at 37°C with 5% CO2, with serum-free ‘complete’ NSC medium. This media consists of DMEM/HAMS-F12 (Sigma D8437) supplemented with N2 and B27 (Life Technologies/Gibco), penicillin, streptomycin (Gibco), BSA (Gibco), b-mercaptoethanol (Gibco), MEM NEAA (Gibco), 1 μg/ml Laminin (Sigma or Cultrex), 10 ng/ml mouse EGF and 10 ng/ml human FGF-2 (Peprotech) Media was exchanged every 3-4 days. Cells were dissociated once 70-80% confluency was reached using Accutase solution (Sigma), passaged approximately 1:6 every 3-4 days. Quiescence was induced by plating cells at a density of 10 cells/mm^2^ in NSC media in the absence of EGF/FGF-2 and supplemented with BMP4 (10 ng/ml, Peprotech). Cells were treated for 1 day or 3 days, as indicated.

### Derivation of genetically engineered cell lines

F6 and F11-19 cell lines were derived previously (Bulstrode *et al*, 2017). Stable transgene integration using the PiggyBac system was used to derive bulk populations of parental ANS4 and FoxO6^-/-^ (53) mouse NSCs with Dox-inducible *FOXG1-V5* overexpression. Cells were transfected using the Amaxa 4D nucleofection system (Lonza) in 16-well cuvette strips, using the DN100 programme. 4 x 10^5^ cells were transfected in 20 μl SG cell line transfection buffer with a total of 800 ng DNA, consisting of the CMV-PiggyBac transposase vector (PBase), pCAG-Tet3G vector (encoding the Tet-On 3G transactivator protein, rtTA) and the TetOn *FOXG1-V5* expression vector in a 2:1:1 ratio. Following recovery, Dox was added (1000 ng/ml) for 24 h. Selection for *FOXG1-V5* expression cassette integration was then commenced by supplementing NSC media with Dox and blasticidin (5 μg/ml). All mock transfected control cells died within seven days of selection. The surviving transfected population were then expanded in NSC media and Dox-inducible *FOXG1-V5* expression was confirmed by ICC and qRT-PCR. Cells were reselected for stable transgene expression between independent experiments. The resulting population were expanded for 3-4 days in NSC media prior to functional assays, during which time existing FOXG1-V5 protein was degraded. Stable transgene integration using the PiggyBac system was also used to derive ANS4 cells with Dox-inducible FoxO6-HA-IRES-mCherry overexpression. The TetOn FoxO6-HA-IRES-mCherry vector was derived using the extensible mammalian modular assembly toolkit (EMMA) system (Martella *et al*, 2017). All EMMA parts are sequence verified, including the FoxO6 coding sequence ordered from GeneArt Gene Synthesis (Thermo Scientific).

For CRISPR/Cas9-mediated gene knockout of *Foxg1* in mouse IENS-GFP, cells were transfected using the Amaxa 4D nucleofection system (Lonza) and the DN100 programme. 1.5 million cells were transfected in 100 μl SG cell line transfection buffer with a total of 4 μg DNA, consisting of 2 μg wild-type Cas9-2A-mCherry vector and 1 μg of each sgRNA plasmid. For sgRNA-encoding plasmids, single-stranded oligonucleotides (IDT) containing the guide sequence of the sgRNAs were annealed, phosphorylated and ligated into BsaI site of U6-BsaI-sgRNA backbone (kindly provided by S. Gerety, Sanger Institute, Cambridge, UK). Three days post-transfection, Cas9-mCherry-expressing cells were isolated by fluorescence-activated cell sorting. Loss of FOXG1 was confirmed and the transfection efficiency estimated in the bulk sorted population by ICC. For derivation of clonal cell lines, 300 cells were plated per 10 cm dish. After 10-15 days, discrete colonies were picked, expanded, and screened for successful disruption of Foxg1 by PCR genotyping and ICC. Loss of FOXG1 protein expression was validated by Western blotting.

CRISPR/Cas9-mediated gene knockout of *FoxO6* in ANS4 cells was performed using a strategy described in (Bressan *et al*, 2017), using two sgRNAs targeting the *FoxO6* exon, Cas9 nickase and a targeting vector comprising an *EF1a*-puromycin antibiotic resistance cassette flanked by 1-kb homology arms specific for the locus. Parental ANS4 cells were transfected using the Amaxa 2B nucleofection system (Lonza). CRISPR/Cas9-mediated HA tagging of FoxO6 was performed using Cas9 RNP ssODN strategy described in (Dewari *et al*, 2018). Once recovered, cells were assessed for successful tag integration by PCR genotyping, ICC and Western blotting.

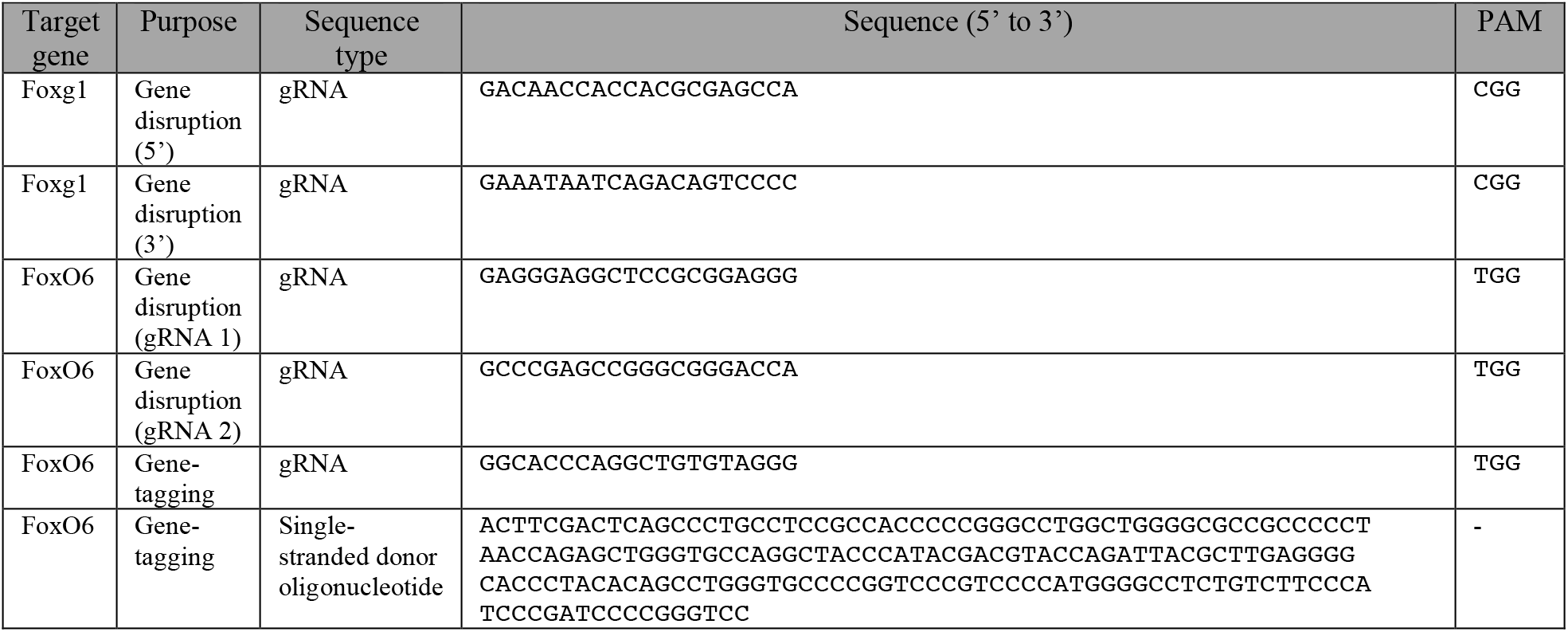

### PCR-based genotyping of genetically engineered cell lines

Genomic DNA (gDNA) isolation from bulk transfected cells and clonal cell lines was performed using the DNeasy Blood and Tissue kit (Qiagen), according to the manufacturer’s protocol. DNA concentrations were quantified using a NanoDrop^TM^ spectrophotometer. All primers were designed using Primer3 software (http://primer3.ut.ee). To identify NHEJ-based indel formation, the region flanking the gRNA target site was amplified using gene-specific primers. In case of *Foxg1* deletion from IENS-GFP, primers were designed flanking the 5’ and 3’ gRNA targeting sites. For validation of FoxO6 gene deletion primers were designed as described in (Bressan *et al*, 2017) (PCR1, 2 and 3). For validation of HA tag knock-in at the *FoxO6* locus, primers were designed flanking the tag, outside of the 77-bp 5’ and 3’ homology arms. PCR products were analysed using 1-2.5% agarose gels with EtBr and GeneRuler™ 1kB plus DNA ladder (Thermo Scientific). Gels were imaged on a UV gel reader or Bio-Rad ChemiDoc™ Imager.

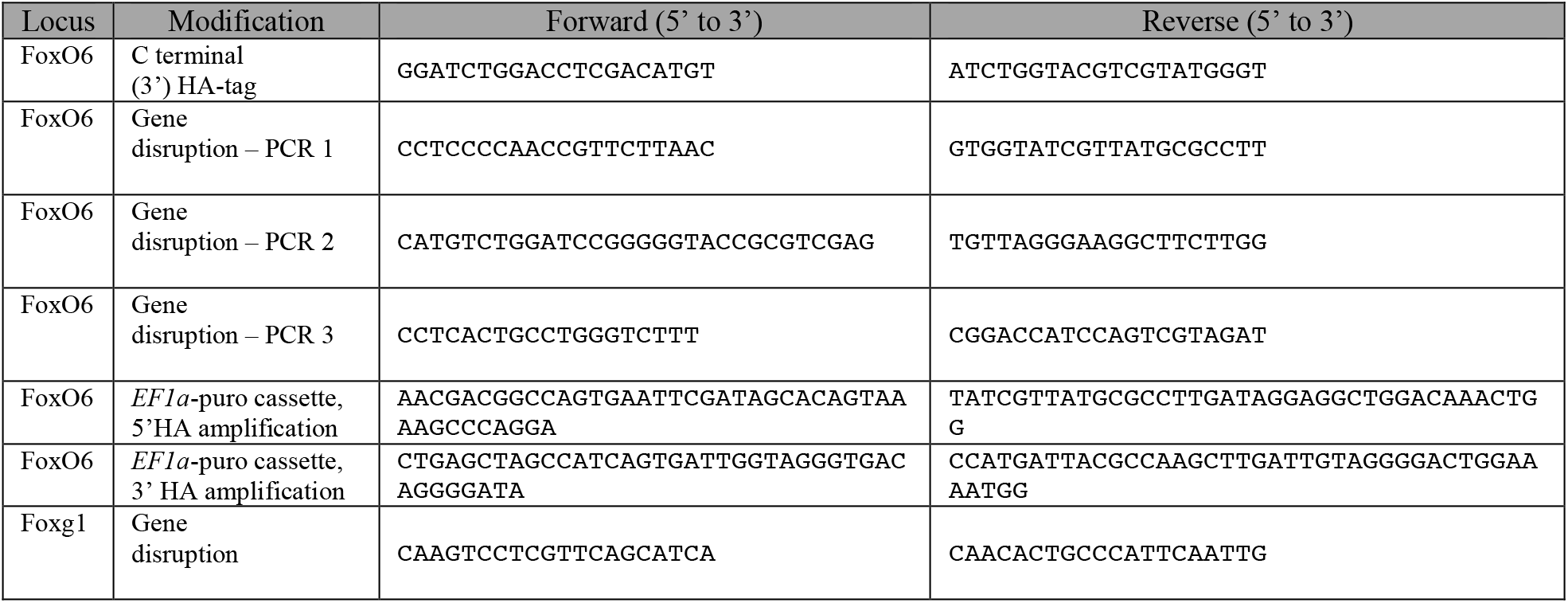

### Immunocytochemistry

Cells were fixed in 4% paraformaldehyde for 10 min, permeabilised in PBS with 0.1% Triton and blocked in 0.1% bovine serum albumin plus 3% goat serum solution for 1 hr at room temperature. Samples were incubated overnight with primary antibodies at 4°C followed by incubation with appropriate secondary antibodies (1:1000; Invitrogen Alexa Fluor™ 488/594/647) for 1 hr at room temperature. Cells were incubated in DAPI (1:10000) for 5 min for nuclear counter-staining. Imaging was performed using the Nikon TiE microscope and NIS software. Analysis was performed using FIJI (Image J) software. Quantification of immunopositive cells was performed using the Cell Counter plugin. Total cell number was determined by DAPI staining. Quantification of FOXG1-V5 staining was performed using PerkinElmer’s Operetta High-content imaging system and Columbus software. The following primary antibodies were used: NESTIN (1:10; Developmental Studies Hybridoma Bank, Rat-401), GFAP (1:1000; Sigma, G3893), FOXG1 (1:100 homemade 17B12 hybridoma), V5 tag (1:2000, eBioscience 14-6796), HA tag (1:100, Cell Signalling Technology 6E2 2367), Ki67 (1:200, ThermoFisher RB-9043-P0), EEA1 (1:200; Cell Signalling Technology 3288) and LAMP1 (1:600; Abcam 25245).

### Western blotting

Immunoblotting was performed using standard protocols. Membranes were blocked in 5% milk in TBS-T (TBS + 0.1% Tween-20) for 1 hr at room temperature and incubated with primary antibody dilutions in 5% milk in TBS-T overnight with rocking. Protein detection was carried out with horseradish peroxidase-coupled secondary antibodies. Membranes were developed using homemade enhanced chemiluminescence (ECL) solution or Clarity ECL Western Blotting substrate (Bio-Rad) and imaged using X ray films or a Bio-Rad ChemiDoc™ Imager. The following primary antibodies were used: V5 tag (1:1000, eBioscience 14-6796), FOXG1 (1:1000, homemade 17B12 hybridoma), GAPDH (1:1000; GenTex, GTX627408), HA tag (1:1000, Cell Signalling Technology 6E2 2367), EEA1 (1:1000; Cell Signalling Technology 3288), LAMP1 (1:1000; Abcam 25245) and Pak1 (1:1000; Cell Signalling Technology 2602). Western blot quantification was performed in FIJI software, normalising to GAPDH loading control.

### Quantitative Real-Time PCR

RNA extraction was performed using the Masterpure^TM^ RNA purification kit according to the manufacturer’s instructions (Epicentre). DNase digestion was performed using RQ1 RNase-free DNase (Promega) or Masterpure^TM^ RNase-free DNase I. RNA concentration was determined using the Qubit™ RNA High Sensitivity kit (Thermo Scientific) or NanoDrop™ Spectrophotometer. Within each experiment, the same amount of RNA was inputted for cDNA synthesis. Reverse transcription was performed using Invitrogen Superscript III. Quantitative RT-PCR (qRT-PCR) was performed using TaqMan Universal PCR Master Mix (Applied Biosystems) and TaqMan gene expression assays (Life Technologies) on a QuantStudio™7 Flex Real-Time PCR machine. No RT and water controls were run on each plate to ensure the absence of contamination. Technical replicates were run to ensure pipetting accuracy. Data were analysed using the ddCt method; this method assumes 100% PCR efficiency which is guaranteed with TaqMan assays. Replicate Ct values were averaged and normalised to the housekeeping gene, Gapdh (to give dCt). These values were then normalised to a calibrator sample (to give ddCt). Data are presented as log_2_(fold change) or - ddCt, where this value equals zero for the calibrator, as indicated in the figure legends. The following TaqMan assays (Life Technologies) were used: hFOXG1 (Hs01850784_s1), mGapdh (Mm99999915_g1), mFoxO6 (Mm00809934_s1), mPlk1 (Mm00440924_g1), mNestin (Mm00450205_m1), mOlig2 (Mm01210556_m1), mAqp4 (Mm00802131_m1), mGfap (Mm01253033_m1), mCdk4 (Mm00726334_s1), mMyc (Mm00487804_m1), mEgfr (Mm00433023_m1), mId1 (Mm00775963), mCd9 (Mm00514275_g1), Pak1 (Mm00817699_m1).

### Cell proliferation assays

Confluence analysis and growth curves were determined using the IncuCyte™ live cell imaging system (Essen Bioscience). Cells were plated at ~25 cells/mm^2^ in NSC media (EGF/FGF-2) in triplicate wells and imaged periodically until confluence was reached. For analysis of proliferation rates, cells were incubated in NSC media (EGF/FGF-2), supplemented with 10 μM EdU for 24 h. Cells were then fixed in 4% PFA for 10 min at room temperature and stained with the Click-iT EdU Alexa Fluor 647 assay kit (Life Technologies) according to manufacturer’s instructions. Imaging was performed using the Nikon TiE microscope and NIS software. For each condition, triplicate wells were analysed (4×4 10x stitched images per well). The total cell number was determined by DAPI staining. Quantification was performed using the Image thresholding and Particle Analysis functions on FIJI software.

### Colony formation assays

Colony formation in NSC media (EGF/FGF-2) was assessed by plating cells at a density of 1 cell/mm^2^ (1000 cells per well of a 6 well plate, with 6 replicate wells). Media was changed every 3-4 days. Following 10 days, plates were fixed using 4% PFA for 10 min at room temperature. Colonies were stained using methylene blue for 30 min. Plates were washed gently with deionised water and allowed to dry. Plates were then imaged on a Celigo™ Image Cytometer (Nexcelom Bioscience). Colonies were counted manually using the Cell Counter plugin on FIJI, or the % pixel area of the well covered by colonies was quantified using FIJI Image thresholding and Particle Analysis functions.

For assessment of colony formation following BMP4 treatment, cells with Dox-inducible FOXG1-V5 overexpression were plated at a density of 10 cells/mm^2^ (10,000 cells per well of a 6 well plate), in NSC media in the absence of EGF/FGF-2 and supplemented with BMP4 (10 ng/ml) (Bulstrode *et al*, 2017). After 24 h, media was replaced fully with NSC media containing EGF/FGF-2 with or without Dox (1000 ng/ml). Media was then replaced every 3-4 days. Following 10-12 days, plates were fixed using 4% PFA for 10 min at room temperature. Colonies were stained using methylene blue for 30 min and imaged on a Celigo™ Image Cytometer (Nexcelom Bioscience). Colonies were counted manually using the Cell Counter plugin on FIJI, or the % pixel area of the well covered by colonies quantified using FIJI Image thresholding and Particle Analysis functions. Three technical replicates were averaged to give the mean number of colonies per biological replicate.

### Imaging analysis of vacuoles

Live imaging following FoxO6-HA induction was performed using the Nikon TiE microscope. Imaging began 4 hours after Dox addition and images were obtained every 10 minutes for ~18 hr. For lysosome assessment, 1000x LysoView488™ stock solution (70057 Biotium) was diluted to 1x in NSC media. Cells were incubated in media containing 1x LysoView488™ for 30 min at 37°C prior to imaging. For lipid droplet assessment, BODIPY 493/503 (Invitrogen D3922, 5 mg/ml) was used. For analysis of Dextran uptake, 70000 MW FITC-Dextran (Invitrogen, 070621, 20 mg/ml) was diluted 1:20 to 1 mg/ml in NSC media and added to cells overnight coincident with Dox addition if appropriate. Dextran uptake was visualised in the green channel by imaging and flow cytometry. EGF uptake was visualised by incubating vacuolated cells with media containing 100 ng/ml of EGFR ligand conjugated a fluorophore (EGF-647, E35351, Thermo Fisher Scientific) for 1 hr prior to washing and imaging.

### Statistical analyses

Statistical analyses were performed in GraphPad Prism 7. Biological replicates were considered as different passage numbers of same cell line plated in independent experiments. Mean and SEM or SD, and n numbers, are shown in the figure legends. Due to small sample sizes, tests for normality and distribution were of limited value. However, this was not considered to be an impediment to parametric analysis with small n numbers (de Winter, 2013). Statistical tests used are indicated in the figure legends. For qRT-PCR data, statistics calculated from ddCt values. Where two-tailed one-sample t-tests are used, this is based on the null hypothesis that log_2_(FC)/-ddCt equals zero (i.e. equal to the calibrator sample). Paired Students t-tests were used where samples (e.g. wild-type and FoxO6 KO cells) must be matched due to variation between biological replicates (e.g. growth analysis, colony assays following BMP4 treatment). Where significant, p values are indicated in Figures as asterisks, * p<0.05, ** p<0.01, *** p<0.001, **** p<0.0001.

## Supplementary Figures

**Figure S1.**
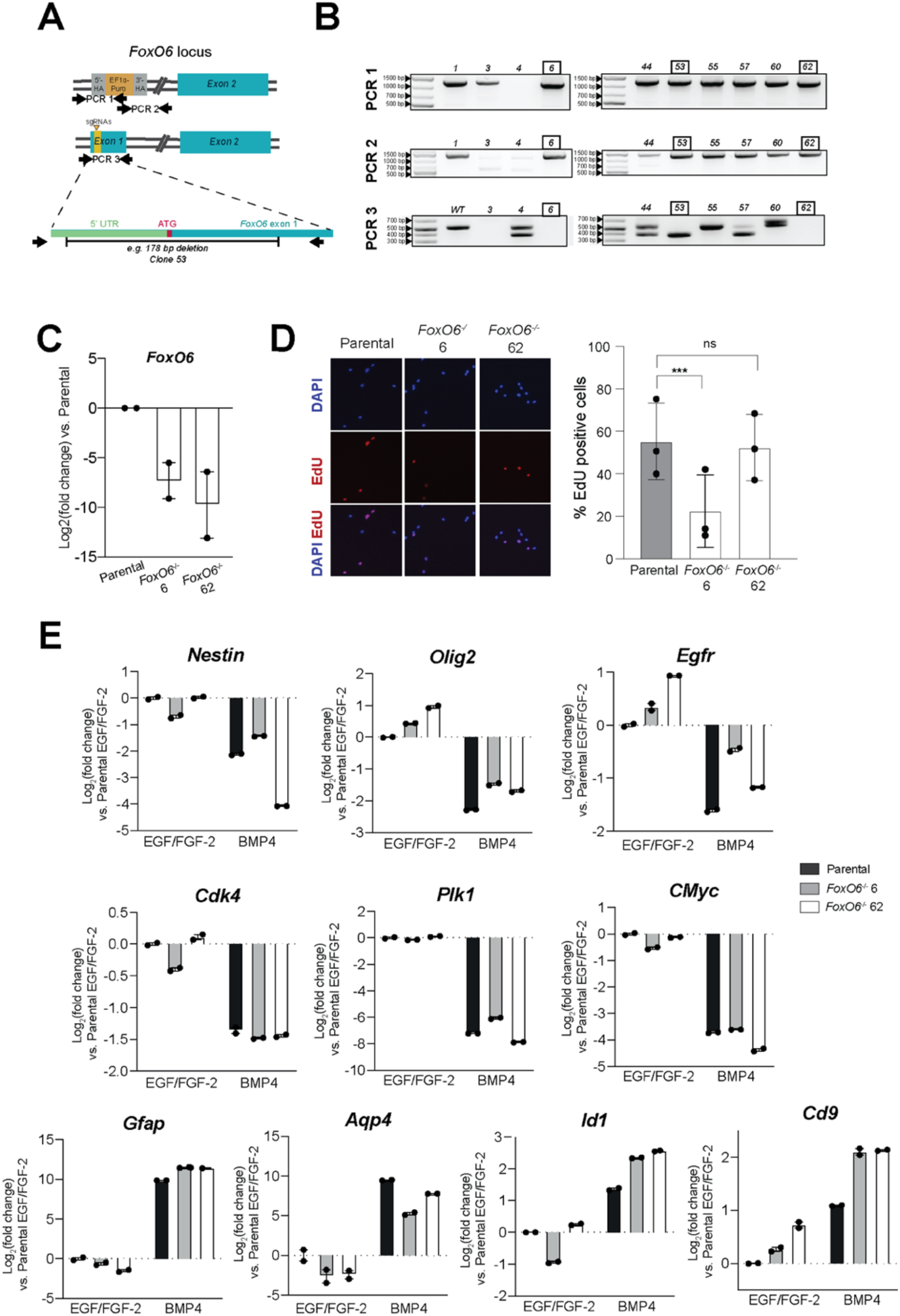
FoxO6 is not essential for NSC proliferation or response to BMP4, related to Figure 3. **(A)** Schematic of *FoxO6* locus following CRISPR/Cas9-mediated knockout strategy. Yellow triangles show the sgRNA target sites, resulting in a 178 bp deletion in allele 1 in FoxO6 KO clone 53. Exon 1 of allele 2 is replaced by an EF1a-puromycin cassette. **(B)** PCR genotyping of *FoxO6* KO clonal cell lines 6, 53 and 62. PCR 1 and PCR 2, across the 5’ and 3’ homology arms of the EF1a-puromycin cassette, respectively, show correct integration at one of the *FoxO6* alleles. PCR 3 shows a 178 bp deletion (53) or loss (6, 62) of the remaining *FoxO6* allele. WT parental band = 565 bp, knockout (53) band = 387 bp. **(C)** qRT-PCR analysis of *FoxO6* mRNA levels in FoxO6 KO clonal cell lines 6 and 62, compared to parental cells (in which log_2_(FC) = 0). Expression values were normalised to Gapdh. Y axis represents log_2_(Fold change). Mean +/− SEM. n=3 independent experiments. Each data point shows the mean of one experiment, performed in technical duplicates. **(D)** EdU incorporation assay (24h pulse) in parental and FoxO6 KO clonal lines (6 and 62) grown in EGF/FGF-2. (Left) Representative fluorescent images of EdU incorporation. (Right) Plot shows mean +/− SEM, n=3 independent experiments. Each data point shows the mean of one experiment performed in technical triplicates. **(E)** qRT-PCR analysis of NSC (*Nestin, Olig2, Egfr*), cell cycle marker (*Cdk4, Plk1, Cmyc*), and astrocyte/quiescence (*Gfap, Aqp4, Id1, Cd9*) marker expression in ANS4 parental and FoxO6 KO clonal cell lines (6 and 62) in EGF/FGF-2 and after 24 h BMP4 treatment. Expression values were normalised to *Gapdh* and shown relative to the expression in parental cells in EGF/FGF-2 (in which log_2_(FC) = 0, shown by the dotted line). Graph shows Mean +/− SD. One experiment, performed in technical duplicates.

**Figure S2.**
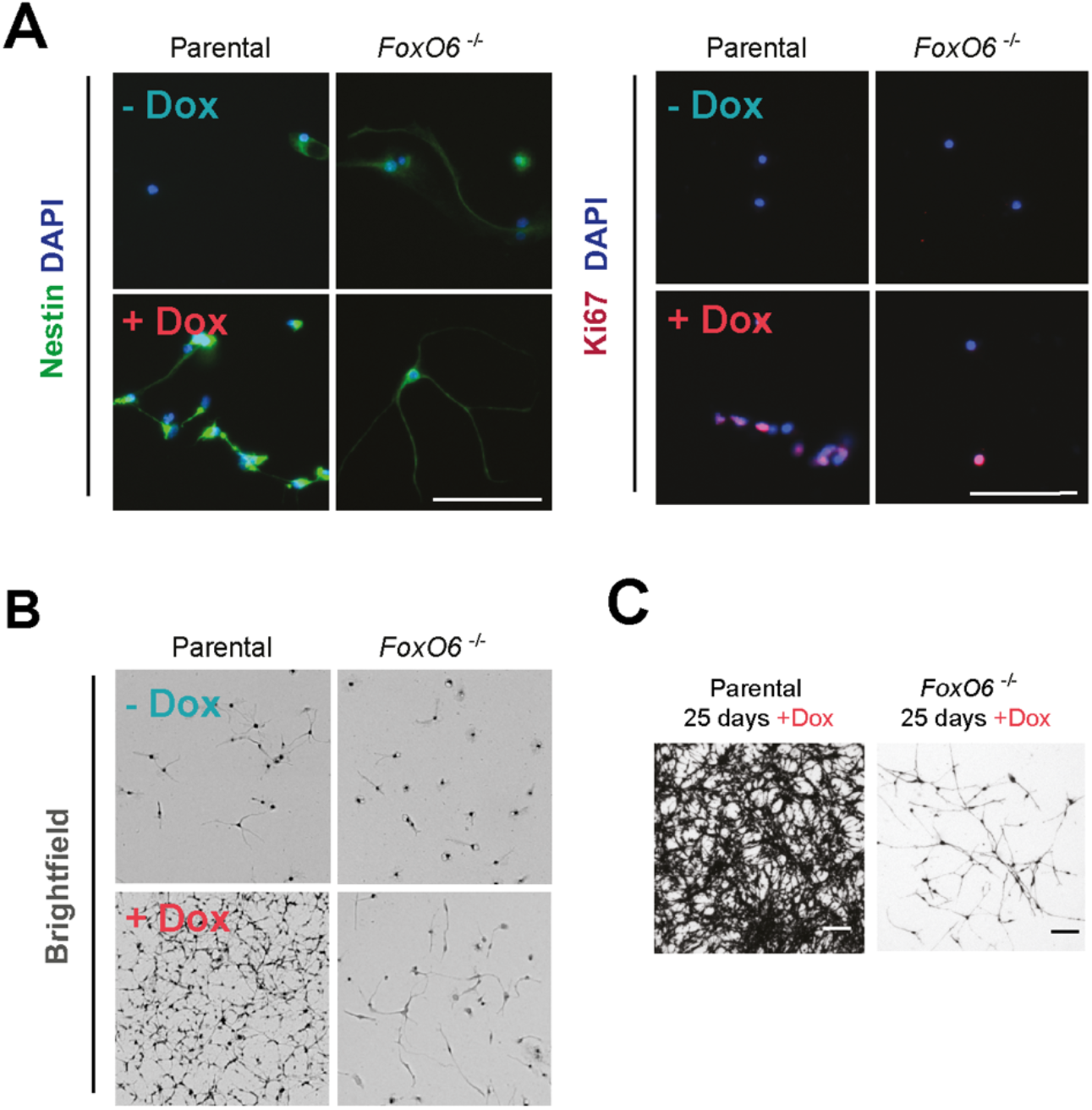
FOXG1-induced reactivation of quiescent NSCs is inhibited in *FoxO6* null cells, related to Figure 4. **(A)** ICC images showing Nestin (left) and Ki67 (right) expression at Day 10 in NSC media with or without Dox addition (following 24 h BMP4 treatment) in both parental and FoxO6^-/-^ 53 cells engineered with inducible FOXG1-V5 construct. **(B)** Representative brightfield images following fixation of colony assay plate and staining with methylene blue. **(C)** Brightfield images of parental and FoxO6 53 colonies after 25 days in NSC media + Dox. Scale bars: 100 μm.

**Figure S3.**
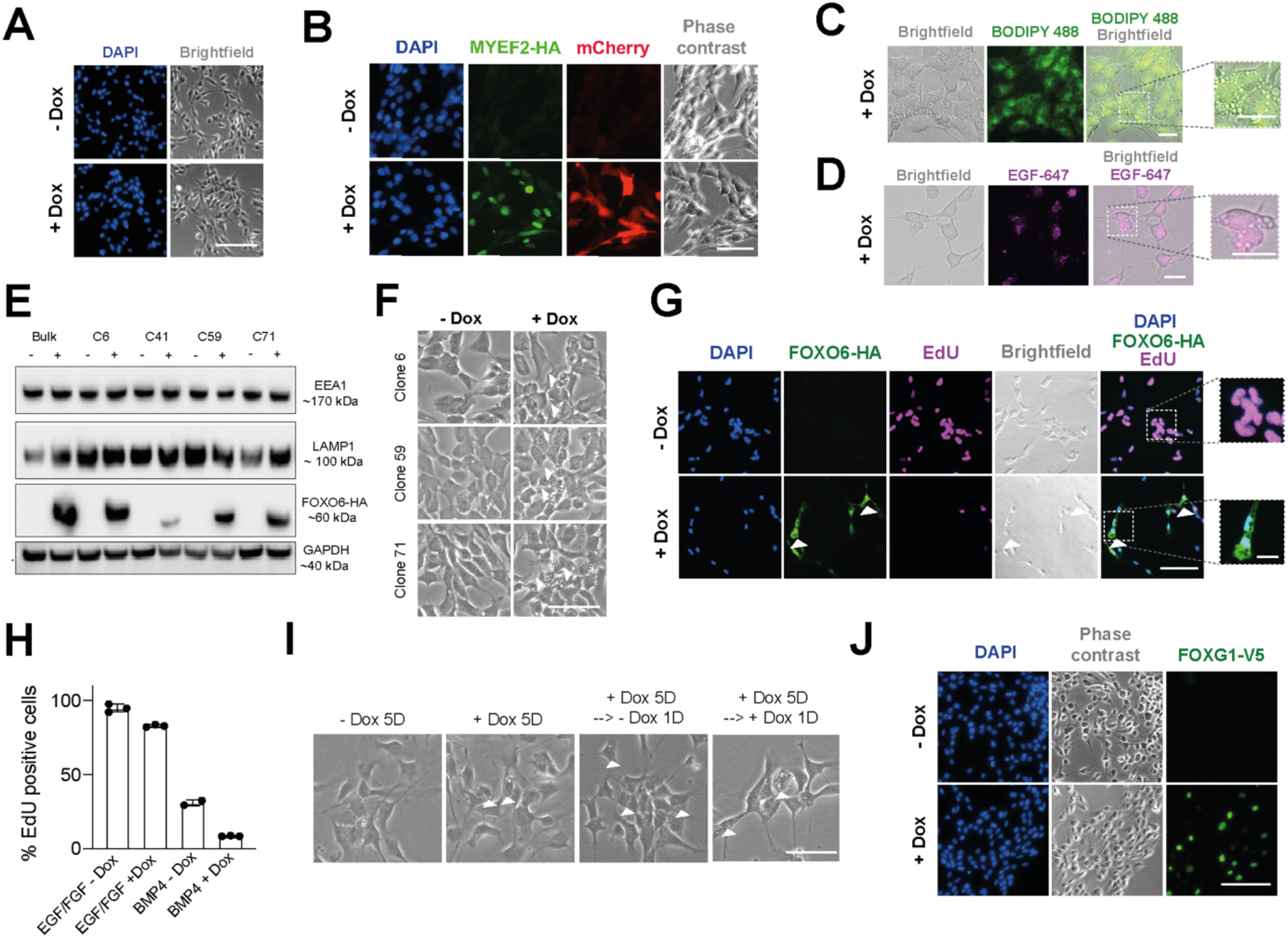
Elevated FoxO6 induces the formation of large acidic vacuoles by macropinocytosis, related to Figure 5. **(A)** Phase contrast imaging following Dox addition to untransfected ANS4 cells does not induce vacuole formation. Scale bar 100 um. **(B)** ICC following Dox-induced (24h) MYEF2-HA-IRES-MCHERRY overexpression in mouse GSC line ‘NPE’ shows no evidence of vacuole formation. Scale bar 50 um. **(C)** BODIPY lipid staining does not colocalise with vacuole structures in FoxO6-inducible cell line (C71, 2 days +/− Dox). Scale bar 25um. **(D)** EGF-647 uptake after a pulse of 1 hr shows puncta representative of receptor-mediated endocytosis (C71 incubated overnight with Dox prior to EGF-647 pulse). Scale bar 25 um. **(E)** Western blot analysis of LAMP1, EEA1 and HA upon FoxO6-HA overexpression (+/− Dox). GAPDH is used as a loading control. Bulk transfected population sorted for mCherry and clonal cell lines (6, 41, 59, 71) analysed. **(F)** Phase contrast images show vacuole formation in FoxO6-HA inducible cell lines following Dox+Dextran overnight incubation, prior to flow cytometry analysis. Scale bar 100 um. **(G)** Imaging of EdU incorporation in FoxO6-HA inducible cells (C71) after 2 days in EGF/FGF +/− Dox (24h pulse). Scale bar 100 um or 25um. **(H)** EdU incorporation after EGF/FGF-2 or BMP4 for 3 days +/− Dox (24h pulse). n=3 technical replicates, mean +/− SD. **(I)** Phase-contrast images of Dox treated cells in culture (c71). Vacuolated cells remain after 5 days in Dox and following Dox removal. **(J)** ICC in F6 cells with Dox-inducible FOXG1-V5 shows no evidence of vacuolisation upon Dox addition (FOXG1-V5 induction). Scale bar 100 um.

